# Extracellular tau clearance is governed by its aggregation state and independent of microglial activation by LPS and IFN-γ

**DOI:** 10.1101/2025.06.23.661190

**Authors:** Andrew Shultz, Anna Vincze, Todd T. Yau, Emma L. Poirier, Hannah Demanche, Jennifer N. Rauch

## Abstract

Microglia are the tissue resident macrophages of the brain and their contribution to tau pathology progression remains to be fully understood. In this study, we developed a quantitative platform to elucidate the processing of extracellular tau within human induced pluripotent stem cell (iPSC)-derived microglia. We show that iPSC-derived microglia internalize monomeric and fibrillar tau through different cellular mechanisms and with different clearance kinetics. Acute inflammatory activation of microglia alters tau endocytosis, but surprisingly does not impact tau clearance. These results highlight the importance of the microglial endo-lysosome system as a regulator of tau pathology that is decoupled from acute microglial activation.

**Highlights:**

- Human iPSC-derived microglia endocytose tau using divergent cellular mechanisms
- NanoBiT system can measure tau endocytosis and degradation in cells
- Aggregation of tau impacts the rate of extracellular clearance after endocytosis
- Acute inflammation affects total endocytosed tau, but not clearance in microglia

## 1. Introduction

Neurodegeneration is one of the prominent pathological hallmarks of Alzheimer’s disease (AD) and numerous non-AD tauopathies. While increasing evidence suggests that pathological modification of the microtubule-associated protein tau and neuroinflammation play a significant role in the neuronal loss observed in tauopathies (Didonna, 2020; Parra Bravo et al., 2024), the exact mechanism by which these two pathologies ultimately contribute to neurodegeneration remain unclear. In disease, tau is known to aggregate into filamentous inclusions and spread throughout the brain in a process that is thought to be mediated primarily through neuronal synapses (Colom-Cadena et al., 2023; Vogel et al., 2020). However, increasing evidence has pointed to the contribution of non-neuronal cell-types, such as microglia, that can also impact tau pathology spread (Asai et al., 2015; Wang et al., 2022).

Microglia serve as the resident immune cells in the central nervous system and have a myriad of roles to maintain overall brain health. Microglia can phagocytose extracellular pathogens, sculpt synapses, and secrete cytokines that reshape the neuronal milieu (Sarlus and Heneka, 2017). Microglia have been highlighted in tauopathy research for their interesting duality serving as both “friend” and “foe” for pathology progression (Odfalk et al., 2022). As professional phagocytes, it has been proposed that microglia may play a helpful role at early stages of disease, helping to internalize and mitigate toxic tau threats (Luo et al., 2015). However, at later disease stages microglia function declines and a breakdown in this protective function occurs that is not well understood. Studies in rodent primary microglia have collectively highlighted that while microglia are capable of internalizing tau aggregates, this process can lead to a hypofunctional state that may actively contribute to tau spread (Brelstaff et al., 2018, 2021; Hopp et al., 2018). Furthermore, ablation of microglia from tauopathy mouse models has been shown to protect against overt neurodegeneration (Shi et al., 2019). Translation of this work to human cells is potentially complicated by species-specific differences in gene expression and microglial functional state (Yvanka De Soysa et al., 2022). Human induced pluripotent stem cell (hiPSC)-derived microglia now offer a tractable platform to interrogate human microglial biology, yet quantitative tools to monitor real-time tau handling in these cells have been limited.

Here we leverage the NanoBiT split-luciferase system (Dixon et al., 2016) to kinetically profile extracellular tau uptake and clearance in cells, including hiPSC-derived microglia. We show that monomeric and aggregated tau are internalized through divergent cellular mechanisms and that this internalized tau is cleared via the lysosome. Crucially, once inside the cell, the aggregation state of tau, rather than the inflammatory status of the microglia, dictates the tempo of turnover, with fibrillar tau persisting significantly longer than its monomeric counterpart. Acute exposure to canonical pro-inflammatory (LPS + IFN-γ) stimuli alters the overall amount of tau internalized but leaves subsequent clearance rates unchanged.

Collectively, our study positions the endo-lysosomal axis—rather than acute inflammatory polarization—as the primary bottleneck for fibrillar tau removal in human microglia. By illuminating how tau aggregation state, but not the cellular activation state, governs intracellular tau processing, we provide a new framework when thinking about the role of microglia in neurodegeneration. We highlight how the microglial contribution to the clearance of pathogenic extracellular tau species is a critical nexus for tau processing that should be considered when developing novel therapeutic strategies.

## 2. Materials and methods

### 2.1. Cell culture models

To produce iTF-Microglia, induced pluripotent stem cells (iPSC) containing a doxycycline inducible expression of six transcription factors (iTF-iPSC) were differentiated following a previously established protocol (Dräger et al., 2022). Briefly, iPSC were dissociated with Accutase and seeded on double coated tissue culture plates (Poly-L-Lysine and Matrigel) with Essential 8 medium, 10 nM ROCK inhibitor and 2 μg/mL doxycycline. On day 2, the media was replaced with Advanced DMEM/F12 containing Antibiotic-Antimycotic (1X), GlutaMAX (1X), 2 μg/mL doxycycline, 100 ng/mL Human IL-34, and 10 ng/mL Human GM-CSF. On day 4, the medium was replaced with the addition of 50 ng/mL Human M-CSF, and 50 ng/mL Human TGFB1. On day 8, iTF-Microglia were either assayed or dissociated with TrypLE Express and plated onto new double coated tissue culture plates (Poly-L-Lysine and Matrigel). All experiments were performed between days 8-15. For CRISPRi, media was replaced every 2-3 days with TMP (20μM).

To produce iMGLs, WTC-CLYBL-dCas9-TagBFP-KRAB-cl391 were differentiated into hematopoietic progenitor cells (HPCs) using a previously published protocol (McQuade et al., 2018). Briefly, iPSC were dissociated with ReLeSR and seeded as aggregates on 12-well Matrigel coated plates with mTeSR™ Plus medium supplemented with 10nM ROCK inhibitor. The next day (Day 0), wells containing 16-40 colonies had their media replaced with Media A from the STEMdiff™ Hematopoietic Kit and a half media change on day 2. On day 3, the media was changed to Media B from the STEMdiff™ Hematopoietic Kit with half media changes on days 5, 7 and 10. On day 12, the floating HPCs were collected and seeded onto Matrigel coated 6 well plates with iMGL differentiation media containing Human IL-34 (100ng/mL), Human M-CSF (25ng/uL) and Human TGFB1 (50ng/uL). Media was added every other day for 25 days and further matured with CD200 (100ng/uL) and CXLC1 (100ng/uL) for 3 days. Microglia were either assayed on day 40 or collected and seeded onto poly-L-Lysine coated plates. All experiments were performed on days 40-46.

To produce iNeurons, WTC11-iPSCs containing a doxycycline inducible expression of Neurogenin2 (Ngn2) were differentiated using a previously published protocol (Chen et al., 2020). Briefly, iPSCs were dissociated with Accutase and seeded on Matrigel coated plates with Knockout DMEM/F12, NEAA (1X), N2 supplement (1X), 10ng/ml NT-3, 10ng/ml BDNF, 1μg/ml mouse laminin and 2 μg/mL doxycycline. Three days later, cells were released with Accutase and seeded on poly-L-Lysine coated plates with DMEM/F12 (0.5X), Neurobasal A (0.5X), NEAA (1X), GlutaMax (0.5X), N2 supplement (1X), B27-VA Supplement (0.5X), 10ng/ml NT-3, 10ng/ml BDNF, 1μg/ml mouse laminin and 2 μg/mL doxycycline. Media was fully replaced on day 3 with the same media without doxycycline. iNeurons were assayed between days 5-11.

H4 cells were cultured in Dulbecco’s Modified Eagle Medium supplemented with 10% FBS and 1% penicillin/streptomycin. Cells were incubated in a humidified incubator with 5% CO2 at 37 °C.

### 2.2. Molecular cloning and stable cell line generation

sgRNA (single guide RNA) plasmids for individual genes were cloned into pLG15 (H4 cells) or pMK1334 (iPSC) vectors following a previously published protocol (Horlbeck et al., 2016). Sequences for sgRNAs used are as follows: NTC sgRNA; GTCCACCCTTATCTAGGCTA, LRP1 sgRNA; GACTTCAGTCCGGGGAACAG, RAB7A sgRNA; GTTGGCCATAAAGCCTGAGG. For Tau-HiBiT constructs, the pRK172 vector containing Tau (2N4R) was digested with BstXI (N-terminus), EcoRI (C-terminus) and PvuII-HF (internal). Using NEB HiFi Assembly, gBlocks containing the HiBiT sequence with GSSG linkers, were assembled into the vector. The correct sequence was confirmed with full plasmid sequencing (Plasmidsaurus).

For lentiviral production, HEK293-T cells were transfected with pMD2.G (3ug), psPAX2 (9ug), sgRNA plasmids (12ug) (PLG15 or pMK1334) using PEI (8ug/mL). Two days after transfection, cell culture media was collected, filtered, and concentrated using Alstem lentivirus precipitation solution. H4i cells (H4 cells containing dCas9-KRAB) and iTF-iPSCs were infected with concentrated sgRNA lentivirus and selected with puromycin (2ug/mL). Knock down was confirmed with RT-qPCR and western blot.

### 2.3. Protein biochemistry

RAP protein was expressed in ClearColi BL21(DE3) following manufacturer recommendations and purified as previously described (Rauch et al., 2020). Tau protein was purified as described (Rauch et al., 2018), protein concentration was measured by BCA. Tau was labeled with Alexa Fluor™ 647 NHS Ester following a previously published protocol (Rauch et al., 2020). Average degree of labeling for tau protein preps was ∼1.

Endotoxin for tau protein preps were removed with a Triton X-114 treatment. Briefly, tau protein was incubated with a final concentration of Triton X-114 of 2% for 1 hour at 4°C. The protein was then incubated in a water bath at 37°C for 10 minutes and centrifuged for 20 minutes, 20,000xg at 37°C. This process was repeated two more times. Protein was passed through a detergent removal spin column to remove residual detergent. Endotoxin concentration was determined by incubating recombinant proteins (Tau and RAP) with HEK-Blue™ hTLR4 Cells overnight. Conditioned medium was collected the next day and was incubated with QUANTI-Blue detection reagent following manufacturer guidelines. Recombinant proteins that were utilized for microglia experiments were used at endotoxin levels <0.1 EU/mL.

For tau fibrillization, tau monomer was incubated with agitation at 37°C with heparin in a 4:1 molar ratio (tau:heparin), DTT (3mM), and 20µM Thioflavin T in DPBS for 3-7 days. Fibrils were isolated after confirmation of ThT fluorescence plateau via ultracentrifugation at 100,000xg for 1 h and resuspended in DPBS. Protein concentration was measured by BCA.

### 2.4. Cellular assays

For tau uptake assay, microglia were incubated with Tau-Alexa Fluor 647 (6.25 to 50nM) for 1 hour (monomer) or 3 hours (fibril) at 37°C – time points were chosen based on our previous studies that require longer incubation times with fibrils to achieve detectable fluorescent signal (Rauch et al., 2018). After incubation, cells were washed 2x with DPBS and incubated with TrypLE™ Express for 10 minutes at 37°C. Cells were then collected and diluted with DPBS and analyzed via flow cytometry on an Attune NxT Flow Cytometer. Propidium iodide was included to exclude non-viable cells. Cells that were not exposed to labeled tau were used to guide gating. To allow comparisons between groups, the positive gate was set so that the positive control (Tau-647 alone) was ∼50%. For RAP competition assays, RAP was added at the same time as tau at a 5-20 molar excess. Specific incubation times, concentrations, and temperatures for each experiment are listed in the figure legends.

For Tau-HiBiT degradation assay, Tau-HiBiT was added to cell cultures at 50-100nM for 1 hour at 37 °C. After incubation, media was removed, and cells were washed with DPBS. Fresh media without Tau-HiBiT was replaced and the cells were left to incubate for various time points up to 6 hours. A sample at t=0 was always taken as a measure of total endocytosis within the 1 hour incubation and for normalization purposes. At various time points (as indicated) media was aspirated, and cells were washed with DPBS. Trypsin-EDTA (0.25%) was added to the cells and incubated for 5 minutes at 37 °C. Trypsin inhibitor (5 mg/mL) was then added to the cells. Cell suspensions were collected and pelleted with centrifugation (500xg). Supernatant was aspirated and cell pellets were resuspended in Promega Nano-Glo® HiBiT Lytic Reagent and processed according to manufacturer recommendations. In some cases, data is presented with raw RLUs that could vary from day-to-day based on number of cells, temperature during luminescent reading, and day-to-day variation in the make-up of fresh NanoBiT lytic reagent. To combine multiple days data was normalized to the average t=0 (100%) and to cells that did not receive Tau-HiBiT protein (0%). For some data, data was normalized to average buffer t=0 (100%) in order to compare the treatment condition to buffer. For LLOMe treatment, H4 cells were treated with 1mM LLOMe for 24 hours before incubation with Tau-HiBiT. For iTF-Microglia stimulation experiments, microglia were pre-treated with LPS (100ng/uL) or interferon-γ (1 µg/mL) before incubation with Tau-HiBiT.

### 2.5. Statistical Analysis

All statistical analyses were performed using GraphPad Prism (Version 10). Where appropriate, two-way analysis of variance (ANOVA) was used to assess the effects of time and treatment, followed by Sidak’s multiple comparisons post-hoc test. One-way ANOVA with Tukey’s post-hoc test was used when comparing more than two groups within a single factor. In experiments with only two groups and a single time point, unpaired two-tailed t-tests were used. Welch’s correction was applied when variances between groups were unequal, as determined by an F-test. In some experiments, data were normalized to the mean t=0 value to account for baseline variability across independent experimental days. This normalization enabled pooling of data and more accurate assessment of relative change over time. In such cases, statistical comparisons were restricted to the relevant post-treatment time point and a t-test was applied. All data points are shown with the mean ± standard deviation (SD) displayed, and the number of biological replicates (n), independent differentiations, statistical tests used, and significance thresholds are indicated in the figure legends.

## 3. Results

### 3.1. Monomeric and fibrillized tau are internalized in iPSC-derived microglia through divergent pathways

The low-density lipoprotein receptor-related protein 1 (LRP1) has been identified as a key regulator of tau endocytosis in neurons and other cell lines (Chen et al., 2024; Cooper et al., 2021; Rauch et al., 2020), however whether LRP1 facilitates tau endocytosis in human microglia has yet to be determined. Thus, to assess whether LRP1 can facilitate microglial endocytosis we adapted our previously established tau endocytosis assay (Rauch et al., 2020) for use in microglial cultures (Figure 1A). Human microglia-like cells were obtained from induced pluripotent stem cells (iPSCs) utilizing a previously established iPSC-line with inducible transcription factor expression (iTF-Microglia) (Dräger et al., 2022). Microglial identity was confirmed with qPCR and immunocytochemistry staining for microglial homeostatic markers (Figures S1A and S1B). Microglia can become inflamed when exposed to lipopolysaccharide (LPS), which is a common contaminant of proteins purified from *E. coli* (Lively and Schlichter, 2018; Monzón-Sandoval et al., 2022; Ye et al., 2020) and recombinant tau preps have high levels of bound LPS due to tau’s lipid binding properties (Barré and Eliezer, 2006). Further, exposure to LPS can modulate microglial phagocytic properties (Parrott et al., 2021). Therefore, to eliminate this potential confounding factor, we developed a protocol for efficient endotoxin removal from recombinant tau preps which resulted in extremely low endotoxin levels (<0.05 EU/mL) (Figure S1C). We labeled our full-length (2N4R) tau protein with Alexa Fluor 647 (“Tau-647”) and incubated monomeric Tau-647 with iTF-Microglial cultures for 1 h at 37°C to promote endocytosis. After incubation, cells were extensively washed and treated with trypsin to digest any cell surface bound Tau-647 before flow cytometry analysis to determine the extent of Tau-647 internalization (Figure 1A). To test the role of LRP1, we introduced a single guide RNA (sgRNA) targeting LRP1 into iTF-Microglia housing an inducible CRISPRi system. Administration of trimethylolpropane (TMP) stabilizes the CRISPRi dCas9-KRAB protein constitutively expressed within these cells and facilitates gene knockdown (Dräger et al., 2022). With this system we achieved a ∼50% knockdown of LRP1 at both the RNA and protein level (Figures S1D and S1E). The iTF-Microglia that received TMP had a significant reduction in Tau-647 positive cells in comparison to their vehicle control counterparts (Figure 1B), consistent with a role for LRP1 in microglial tau uptake. As an additional control, we confirmed no change in transferrin endocytosis when LRP1 was knocked down (Figure S1F).

**Figure 1.**
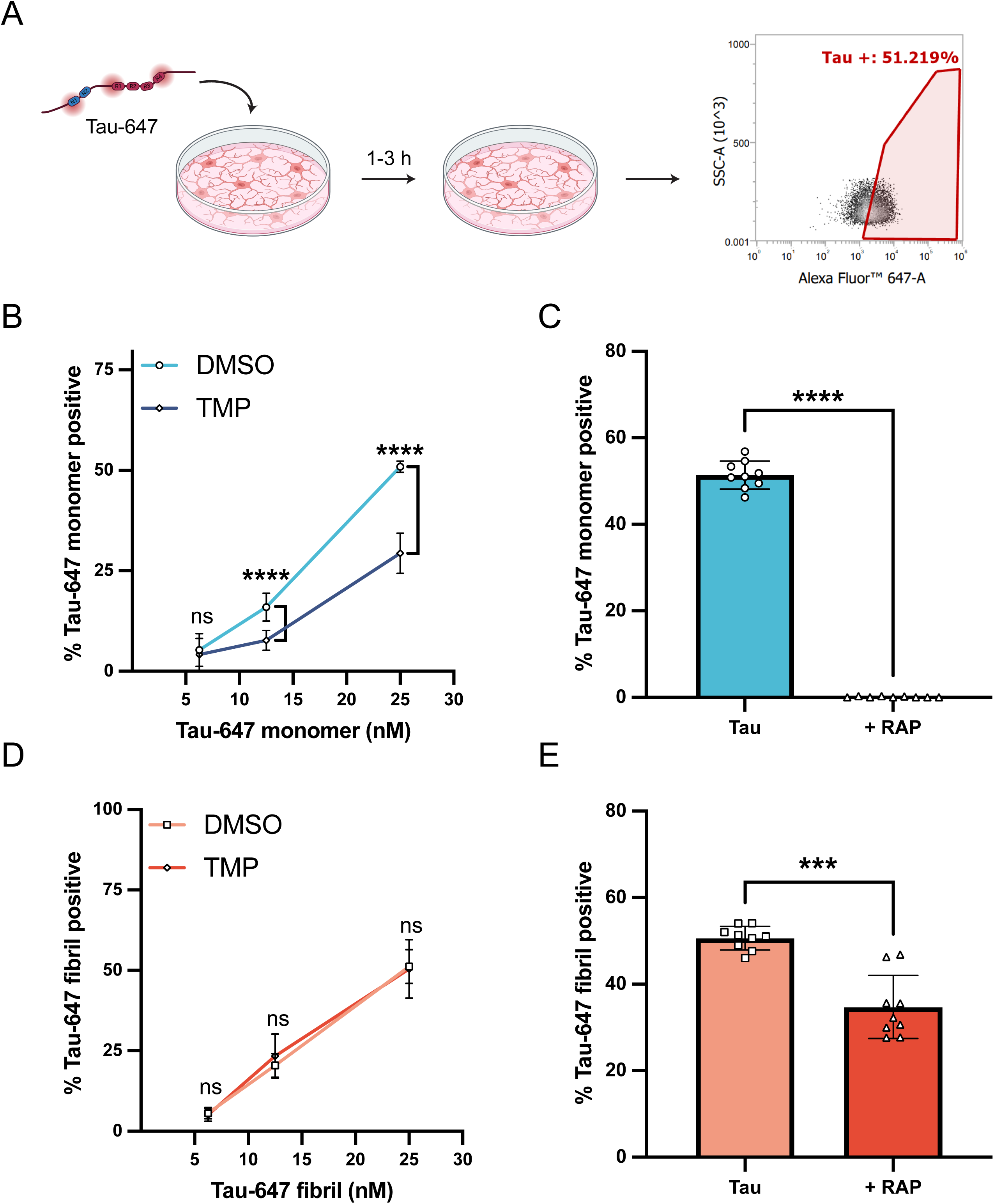
LRP1 facilitates monomeric tau endocytosis in iPSC-derived microglia. (A) Diagram of tau uptake assay. Tau-647 is added to microglial cultures and incubated for 1-3 h at 37°C to allow for internalization. Cells are washed, treated with TrypLE Express enzyme, and analyzed for fluorescence via flow cytometry. (B) Monomeric tau uptake in iTF-Microglia with (TMP, 20µM) or without (DMSO) LRP1 knockdown (mean±SD; n ≥ 6 from three independent differentiations; ns = not significant, ∗∗∗∗p < 0.0001 using a two-way ANOVA; concentration, *F*(2, 42) = 446.7, *p* < 0.0001; cell-line, *F*(1, 42) = 97.77, *p* < 0.0001; interaction, *F*(2, 42) = 32.80, *p* < 0.0001; Sidak’s multiple comparisons test, p values indicated on figure). (C) Tau uptake in iTF-Microglia incubated with monomeric Tau-647 (50nM) or Tau-647 (50nM) + RAP (500nM) for 1 h at 37°C (mean±SD; n = 9 from three independent differentiations; ∗∗∗∗p < 0.0001 using unpaired Welch’s t-test). (D) Fibrillized tau uptake in iTF-Microglia with (TMP, 20µM) or without (DMSO) LRP1 knockdown (mean±SD; n ≥ 6 from three independent differentiations; ns = not significant using a two-way ANOVA; concentration, *F*(2, 42) = 241.1, *p* < 0.0001; cell line, *F*(1, 42) = 0.1125, *p* = 0.7390; interaction, *F*(2, 42) = 0.5470, *p* = 0.5827; Sidak’s multiple comparisons test, p values indicated on figure). (E) Tau uptake in iTF-Microglia incubated with fibrillized Tau-647 (50nM) or Tau-647 (50nM) + RAP (1µM) for 3 h at 37°C (mean±SD; n = 9 from three independent differentiations; ∗∗∗p = 0.0001 using unpaired Welch’s t-test).

Our LRP1 sgRNA iTF-Microglia achieved only a partial reduction in LRP1 levels, thus we rationalized a more complete inhibition of LRP1 may produce a stronger phenotype in our system. We set out to further confirm the relevance of LRP1 in human microglial tau endocytosis by performing the same endocytosis experiment in the presence of excess RAP, a well-characterized LRP1 chaperone/ligand (Bu and Rennke, 1996; Herz et al., 1991). When Tau-647 was co-incubated with RAP, there was a drastic decrease in tau internalization, suggesting a role for LRP1 in microglial tau endocytosis (Figure 1C). We confirmed these findings in an additional iPSC-derived microglia line utilizing a longer 40-day differentiation protocol (iMGLs) (McQuade et al., 2018) (Figure S1G).

To assess whether LRP1 could also facilitate the endocytosis of larger tau aggregates in microglia, we repeated our tau uptake experiments with fibrillized Tau-647 protein (Figures S1H-J). In contrast to our results with monomeric tau, LRP1 CRISPRi knockdown did not change fibrillized Tau-647 endocytosis (Figure 1D). To further inhibit LRP1 we also performed uptake of fibrillized Tau-647 with excess RAP and found a decrease in endocytosis (Figure 1E and Figure S1L: average 51 ± 3% Tau-647 positive to average 35 ± 7% Tau-647 positive) that was significant although less striking as compared to the results we observed with monomeric Tau-647 (Figure 1C and Figures S1K: average 51 ± 3% Tau-647 positive to average 0.1 ± 0.1% Tau-647 positive). Taken together, we conclude that LRP1 is critical for human microglia monomeric tau endocytosis but that fibrillized tau can likely utilize multiple mechanisms for endocytosis, some of which are independent of LRP1.

### 3.2. Development of the NanoBiT system to monitor tau lysosomal degradation

Following endocytosis, proteins are typically trafficked through the endosomal system for eventual degradation in the lysosome. While the use of fluorescently labeled tau is a convenient and high-throughput method to investigate tau endocytosis, it is not a suitable method to measure tau lysosomal degradation as fluorescent dyes have been shown to be longer lived than the proteins they are conjugated to (Mandrekar et al., 2009). Therefore, to investigate the kinetics of tau degradation after endocytosis, we established a cellular tau degradation assay based on the principles of the NanoLuc binary technology system (NanoBiT) (Dixon et al., 2016). In the NanoBiT system, the Nanoluciferase enzyme has been split into a larger 18-kDa subunit (LgBiT) and an 11-amino acid high-affinity peptide (HiBiT) that upon complementation in the presence of substrate produces luminescence. We appended the HiBiT sequence onto 2N4R tau with a short GSSG linker, hereafter referred to as Tau-HiBiT. We created three Tau-HiBiT constructs, with the HiBiT sequence localized to the N-terminus, C-terminus, or internally – at the end of the 2N domain (Figure 2A). Irrespective of the HiBiT location, we were able to measure Tau-HiBiT concentrations from 100nM down to 1pM *in vitro* and concluded that all three Tau-HiBiT proteins were capable of complementing LgBiT to comparable levels (Figure 2C).

**Figure 2.**
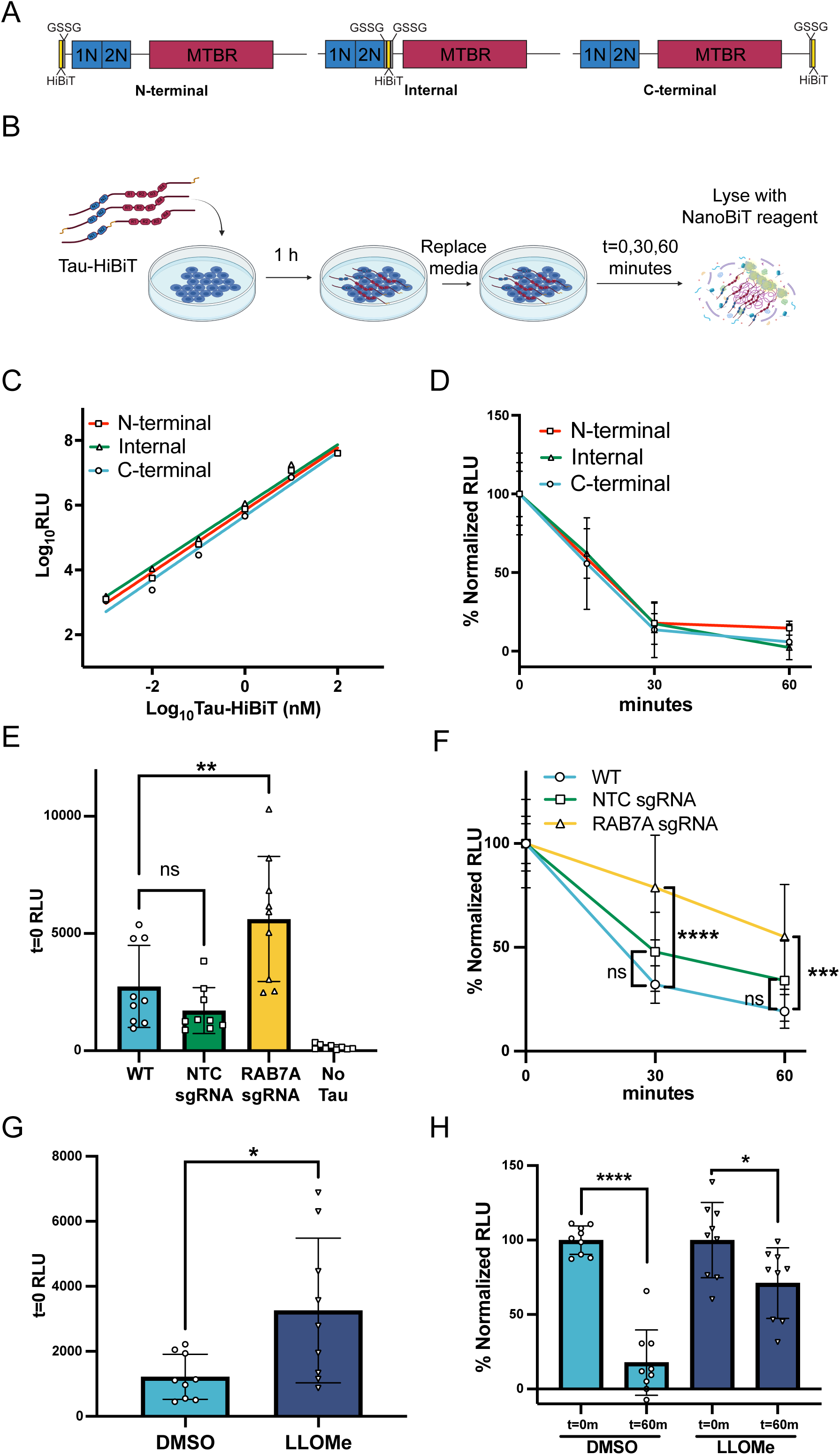
Internalized tau is degraded by the lysosome as measured using the NanoBiT system. (A) Tau (2N4R isoform)-HiBiT constructs for degradation experiments. HiBiT sequence is placed N-terminal, Internal or C-terminal with a GSSG linker. 1N, 2N; N-terminal inserts, MTBR; microtubule binding repeats. (B) Schematic illustrating the Tau-HiBiT clearance assay workflow. Tau-HiBiT is added to cell cultures and incubated for 1 h at 37°C to allow for internalization. Cells are washed, media-free of Tau-HiBiT is added, cells are lifted with trypsin at various time points and analyzed for luminescence using the Nano-Glo® HiBiT Lytic Reagent containing LgBiT and luminescent substrate. (C) Luminescence of Tau-HiBiT proteins (1pM to 100nM) (mean±SD; n = 2). (D) Tau-HiBiT degradation assay of all three Tau-HiBiT constructs (100nM) in H4 cells, samples were normalized to respective initial uptake (t=0) (mean±SD; n ≥ 6 independent cultures). (E) Initial uptake (t=0) of Tau-HiBiT in wild-type (WT), Non-targeting control (NTC) sgRNA, or RAB7A sgRNA H4 cells as measured by luminescence at t=0 (mean±SD; n = 9 independent cultures; ns = not significant, ∗∗ p < 0.01 using a one-way ANOVA, *F*(3, 32) = 17.10, *p* < 0.0001; Tukey’s multiple comparisons test, with p values indicated on figure). (F) Degradation of Tau-HiBiT in wild-type (WT), Non-targeting control (NTC) sgRNA, or RAB7A sgRNA H4 cells. Luminescent signal is normalized to respective initial uptake levels (100%) and cells that were not exposed to Tau-HiBiT (0%) (mean±SD; n = 9 independent cultures; ns = not significant, ∗∗∗ p < 0.001 using a two-way ANOVA; time: *F*(1, 48) = 10.63, *p* = 0.0021; cell line: *F*(2, 48) = 21.73, *p* < 0.0001; interaction: *F*(2, 48) = 0.4505, *p* = 0.6400; Sidak’s multiple comparisons test, p values indicated on figure; t=0 values are shown for reference but were excluded from statistical analysis). (G) Initial uptake (t=0) of Tau-HiBiT in H4 cells pre-treated with vehicle (DMSO) or LLOMe (1mM) for 24 h (mean±SD; n = 9 independent cultures; ∗ p < 0.05 using unpaired Welch’s t-test). (H) Degradation of Tau-HiBiT in H4 cells pre-treated with vehicle (DMSO) or LLOMe (1mM) for 24 h. Luminescent signal is normalized to respective initial uptake levels (100%) and cells that were not exposed to Tau-HiBiT (0%) (mean±SD; n = 9 independent cultures; ns = not significant, ∗∗∗∗ p < 0.0001, ∗ p < 0.05 using unpaired Welch’s t-test).

We next brought the NanoBiT system into cells by developing a Tau-HiBiT degradation assay initially using an H4 neuroglioma cell line. In the Tau-HiBiT degradation assay H4 cells are incubated with Tau-HiBiT (50-100nM) for 1 h to facilitate endocytosis, cells are then washed, and media free of Tau-HiBiT is added to the cultures for various “clearance phase” times. Cells are collected at various times post media change, briefly treated with trypsin to remove any cell surface-bound Tau-HiBiT, and then lysed open with a lysis buffer containing LgBiT and substrate (Figure 2B). To determine the rate of degradation, the luminescence signal is normalized to replicate wells that were incubated with Tau-HiBiT and were immediately lysed (t=0) as this provides a measure of the initial internalization (100%) and to replicate wells that were not exposed to Tau-HiBiT (0%). We have found that initial RLU (t=0) can vary day-to-day based on number of cells, temperature during luminescent reading, and due to fact that the NanoBiT lytic reagent is made fresh every day. Therefore, a normalization of individual days to t=0 is necessary for combining data sets from multiple days and for making rigorous conclusions between cell lines and conditions. After 60 minutes of clearance phase, almost no luminescence over the background could be measured (Figure 2D), suggesting complete degradation of endocytosed Tau-HiBiT within this time frame. All monomeric Tau-HiBiT constructs displayed similar degradation kinetics, so we elected to only use the C-terminus Tau-HiBiT for future experiments.

To confirm that the reduction of luminescence over time is attributed to Tau-HiBiT being proteolytically cleaved within the lysosome we performed both genetic and pharmaceutical manipulations. For the genetic approach, we used CRISPRi to knockdown RAB7A expression in H4 cells (Figure S2A and S2B). RAB7A is a known regulator of late endosome-lysosome fusion (Néel et al., 2024), thus RAB7A knockdown is predicted to prevent tau degradation in our platform. Consistent with this hypothesis, knockdown of RAB7A caused an increase in luminescence at t=0 suggesting higher initial steady state tau levels (Figure 2E). When normalized to this initial higher level, Tau-HiBiT also showed slowed degradation in RAB7A knockdown cells compared to wild-type cells or non-targeting control (NTC) sgRNA cells (Figure 2F). We further confirmed that our platform was reliant on the lysosome using a complementary chemical approach with the small molecule L-Leucyl-L-Leucine methyl ester (LLOMe). LLOMe is a cathepsin C substrate that when cleaved permeabilizes lysosomal membranes and promotes lysosomal leakage into the cytosol, effectively disrupting lysosomal activity (Kavčič et al., 2020; Repnik et al., 2017). Consistent with our RAB7A results, H4 cells receiving LLOMe treatment had an increase of luminescence at t=0 (Figure 2G), suggesting higher initial steady state tau levels. Additionally, cells receiving LLOMe showed a reduced rate of tau degradation with a less significant decrease in luminescence after a 1 h clearance phase (Figure 2H). With these data, we conclude that the NanoBiT system provides a robust platform for measuring endocytosed tau degradation kinetics and that internalized tau degradation is reliant on the lysosomal system.

### 3.3. Modifications to tau structure differentially impact degradation kinetics

Alterations to tau protein structure is a shared characteristic of a variety of neurodegenerative diseases. Point mutations to tau protein’s primary sequence is causative for familial forms of Frontotemporal Dementia (FTD) (Delisle et al., 1999; Iijima et al., 1999) and increases in tau protein post-translational modifications are linked to a swath of tauopathies, including FTD and AD (Wesseling et al., 2020). Therefore, to better understand how changes to tau protein structure can impact degradation kinetics in our model, we designed a series of experiments to test the hypothesis that disease-associated modifications (mutation, phosphorylation, and aggregation) could slow tau degradation within cells.

The FTD tau mutation N279K has been shown to have a resistance to lysosomal proteolytic cleavage *in vitro* (Sampognaro et al., 2023). We therefore produced a TauN279K-HiBiT protein and tested if it had a resistance to degradation in our cellular system. We found that the TauN279K-HiBiT degradation rate had no significant difference from the wild-type (WT) Tau-HiBiT degradation rate (Figure 3A). Next, we asked if tau phosphorylation could impact degradation rate, as tau hyperphosphorylation is a known corollary with disease. The GSK3β kinase is known to phosphorylate tau and GSK3β dysregulation has been hypothesized to contribute to disease progression (Chakraborty et al., 2024; Rankin et al., 2007; Wegmann et al., 2021). Therefore, we phosphorylated Tau-HiBiT *in vitro* using recombinant GSK3β to produce a phosphorylated Tau-HiBiT (“pTau-HiBiT”) (Figure S3A). We found that the pTau-HiBiT degradation rate had no significant difference from the WT Tau-HiBiT degradation rate (Figure 3B).

**Figure 3.**
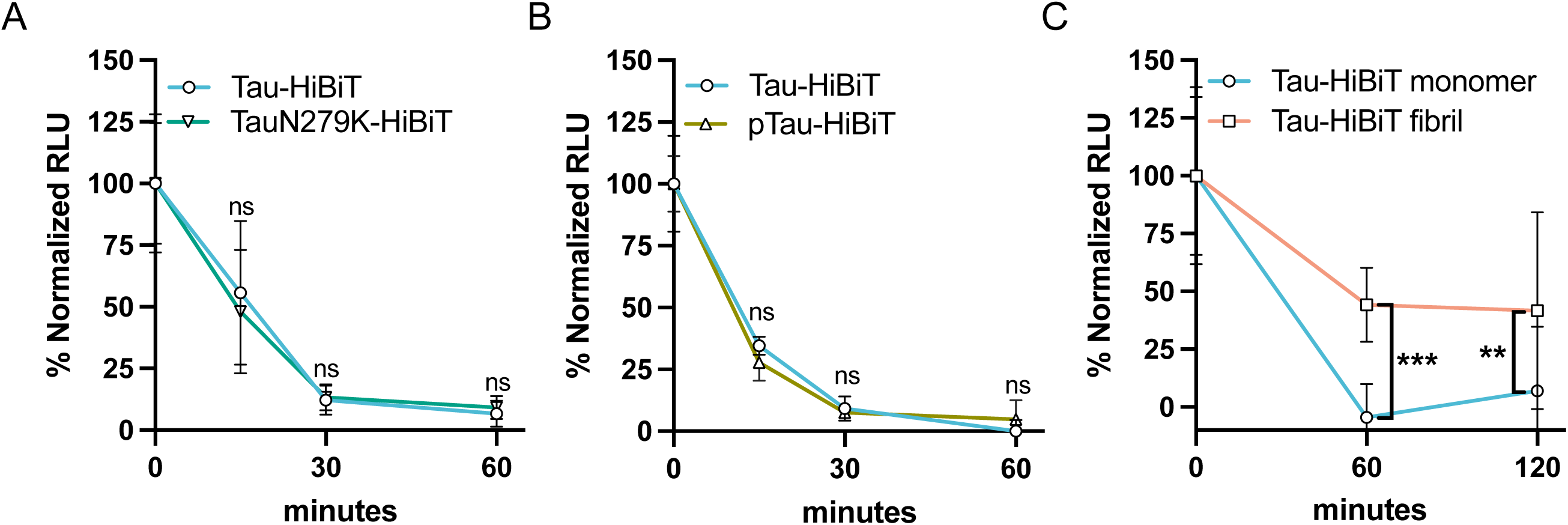
Fibrillization impacts Tau-HiBiT degradation. (A) Tau-HiBiT degradation assay of Tau-HiBiT or TauN279K-HiBiT (100nM) in H4 cells, samples were normalized to respective initial uptake (t=0; 100%) and cells that were not exposed to Tau-HiBiT (0%) (mean±SD; n ≥ 6 independent cultures; ns = not significant using a two-way ANOVA; time: *F*(2, 36) = 26.06, *p* < 0.0001; construct: *F*(1, 36) = 0.05494, *p* = 0.8160; interaction: *F*(2, 36) = 0.3343, *p* = 0.7180; Sidak’s multiple comparisons test, *p* values indicated on figure, t=0 values are shown for reference but were excluded from statistical analysis). (B) Tau-HiBiT degradation assay of Tau-HiBiT or pTau-HiBiT (100nM) in H4 cells, samples were normalized to respective initial uptake (t=0; 100%) and cells that were not exposed to Tau-HiBiT (0%) (mean±SD; n = 6 independent cultures; ns = not significant using a two-way ANOVA; time: *F*(2, 30) = 94.36, *p* < 0.0001; construct: *F*(1, 30) = 0.4521, *p* = 0.5065; interaction: *F*(2, 30) = 3.345, *p* = 0.0488; Sidak’s multiple comparisons test, *p* values indicated on figure, t=0 values are shown for reference but were excluded from statistical analysis). (C) Tau-HiBiT degradation assay of Tau-HiBiT monomer or Tau-HiBiT fibril (70nM) in H4 cells, samples were normalized to respective initial uptake (t=0; 100%) and cells that were not exposed to Tau-HiBiT (0%) (mean±SD; n = 12 replicates from three independent cultures; ∗∗ p < 0.01, ∗∗∗ p < 0.001 using a two-way ANOVA; time: *F*(1, 43) = 0.2958, *p* = 0.5893; protein: *F*(1, 43) = 26.29, *p* < 0.0001; interaction: *F*(1, 43) = 0.7387, *p* = 0.3949; Sidak’s multiple comparisons test, *p* values indicated on figure, t=0 values are shown for reference but were excluded from statistical analysis).

Lastly to understand if aggregation of tau can impact degradation rate, we fibrillized our Tau-HiBiT protein and confirmed that it was still capable of NanoLuc complementation *in vitro* (Figure S3B). We then measured degradation of fibril Tau-HiBiT and found a drastic decrease in the rate of degradation, indicated by a significant difference in the amount of Tau-HiBiT still present after 60 and 120 minutes of clearance phase (Figure 3C). Taken together, our results suggest that while mutation (N279K) and phosphorylation do not have a measurable impact in our cellular tau degradation assay, fibrillization has a noticeable impact on the ability for cells to clear endocytosed tau.

### 3.4. Human iPSC-derived microglia and iPSC-derived neurons have diverse clearance rates

Our initial development of the cellular tau degradation assay was performed in an immortalized neuroglioma cell line (H4). To examine tau processing in a more physiological context, we applied our Tau-HiBiT clearance assays to both monomeric and aggregated tau in iPSC-derived microglia (iTF-Microglia) and iPSC-derived neurons (iNeurons) (Chen et al., 2020). Using iTF-Microglia, monomeric Tau-HiBiT was rapidly depleted after 1 h of clearance phase (Figure 4A). In contrast, aggregated Tau-HiBiT luminescence persisted for up to at least 4 h of clearance phase with 20 ± 10% still remaining (Figure 4A). When testing the frontotemporal dementia–linked TauN279K variant, we observed a modest but statistically significant resistance to clearance compared to wild-type Tau-HiBiT (Figure 4B). Interestingly, this effect was not detected in H4 cells, suggesting that the impact of this mutation may be subtle and cell type–specific, potentially compounding over time in disease. In iNeurons, clearance kinetics were much slower overall—both for monomeric Tau-HiBiT (Figure 4C) and for fibrillar Tau-HiBiT (Figure 4D)—than in iTF-Microglia. Taken together, these findings indicate that, as in H4 cells, iPSC-derived microglia display distinct capacities to process monomeric versus fibrillar tau, possibly reflecting differences in internalization/endocytic routing (Figure 1). Furthermore, iNeurons exhibit a markedly reduced ability to clear extracellular tau compared to microglia, consistent with the accumulation of tau aggregates in neurons in tauopathies.

**Figure 4.**
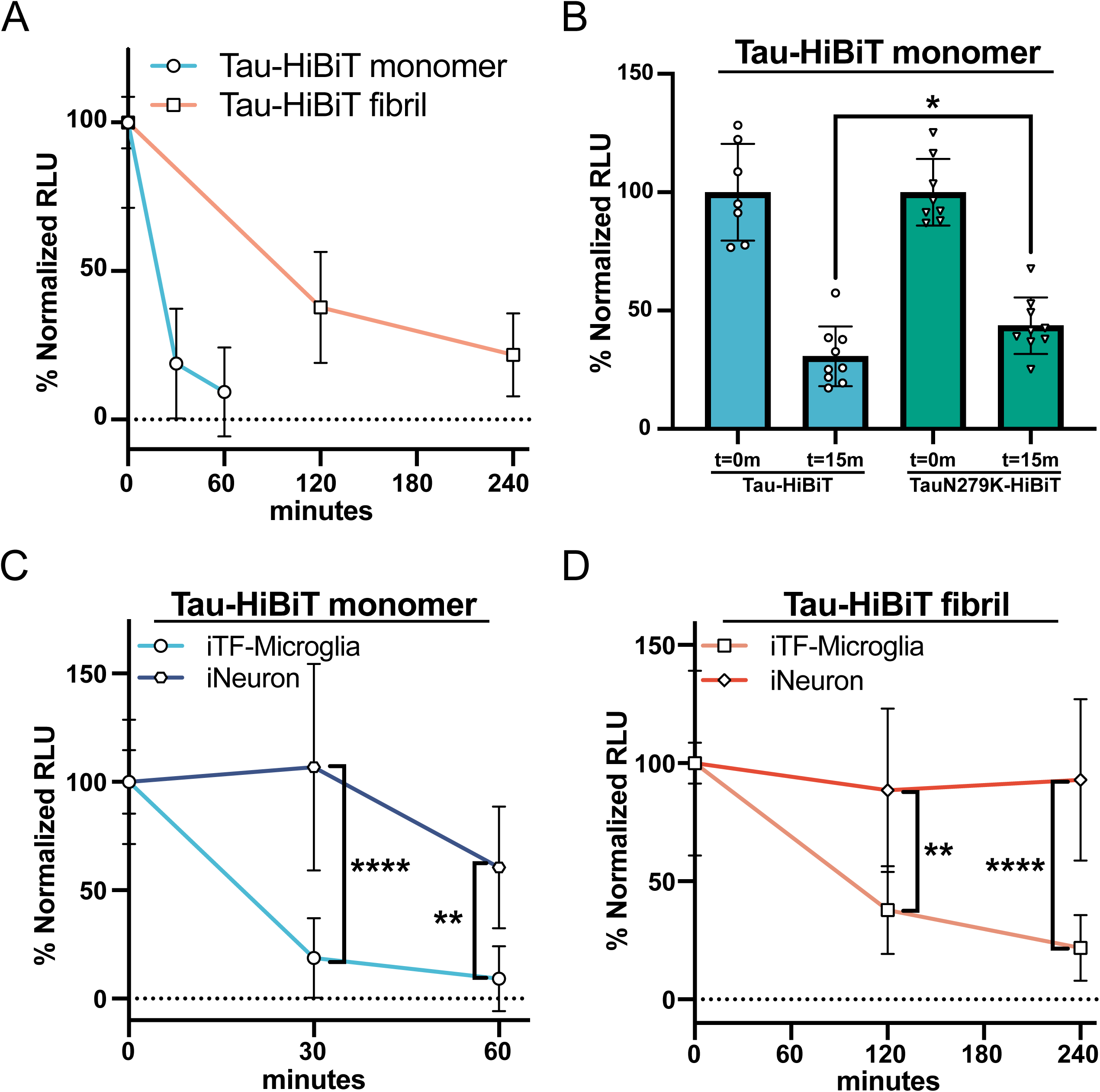
iTF-Microglia and iNeurons show differing abilities to clear endocytosed Tau-HiBiT. (A) iTF-Microglia were subjected to the Tau-HiBiT clearance assay with monomeric Tau-HiBiT (100nM) or fibrillar Tau-HiBiT (50nM), samples were normalized to respective initial uptake (t=0; 100%) and cells that were not exposed to Tau-HiBiT (0%) (mean±SD; n = 9 from three independent differentiations). (B) iTF-Microglia were incubated with monomeric Tau-HiBiT or TauN279K-HiBiT (100nM), samples were normalized to respective initial uptake (t=0; 100%) and cells that were not exposed to Tau-HiBiT (0%) (mean±SD; n ≥ 7 from three independent differentiations; ∗ p < 0.05 using unpaired Welch’s t-test). (C) iTF-Microglia or iNeurons were incubated with monomeric Tau-HiBiT (100nM), samples were normalized to respective initial uptake (t=0; 100%) and cells that were not exposed to Tau-HiBiT (0%) (mean±SD; n ≥ 6 from two-three independent differentiations; ∗∗ p < 0.01, ∗∗∗∗ p < 0.0001 using a two-way ANOVA; time: *F*(1, 28) = 7.489, *p* = 0.0107; cell type: *F*(1, 28) = 46.61, *p* < 0.0001; interaction: *F*(1, 28) = 3.229, *p* = 0.0831; Sidak’s multiple comparisons test, *p* values indicated on figure, t=0 values are shown for reference but were excluded from statistical analysis). (D) iTF-Microglia or iNeurons were incubated with fibrillar Tau-HiBiT (50nM), samples were normalized to respective initial uptake (t=0; 100%) and cells that were not exposed to Tau-HiBiT (0%) (mean±SD; n ≥ 6 from two-three independent differentiations; ∗∗ p < 0.01, ∗∗∗∗ p < 0.0001 using a two-way ANOVA; time: *F*(1, 26) = 0.3888, *p* = 0.5384; cell type: *F*(1, 26) = 43.09, *p* < 0.0001; interaction: *F*(1, 26) = 1.209, *p* = 0.2816; Sidak’s multiple comparisons test, *p* values indicated on figure, t=0 values are shown for reference but were excluded from statistical analysis).

### 3.5. LPS and IFN-γ modulate tau endocytosis but not clearance rate in human iPSC-derived microglia

It is well established that microglia are a highly plastic and versatile cell type that react and respond to a plethora of extracellular stimuli (Lively and Schlichter, 2018; Rock et al., 2005; Woodburn et al., 2021), and recent advances in single-cell transcriptomics have characterized the heterogeneity of microglia states (Martins-Ferreira et al., 2025). Therefore, we hypothesized that stimulation of our iTF-Microglia to an inflamed state could modulate their ability to clear Tau-HiBiT. To test this hypothesis, we first stimulated our iTF-Microglia with lipopolysaccharide (LPS), a known pro-inflammatory driver in microglia (Hoogland et al., 2015). iTF-Microglia cultures were incubated with LPS (100ng/mL) for 24 hours, conditioned media was collected and analyzed with a cytokine array. LPS-stimulated iTF-Microglia media had an increase of proinflammatory cytokines such as IL-6 and IL-8 (Figure S4A and S4B), confirming that our iTF-Microglia were responding to the LPS treatment. We then tested whether LPS stimulation altered endocytosis or clearance kinetics of endocytosed Tau-HiBiT. We found that when iTF-Microglia were primed with LPS for 24 hours and then incubated with Tau-HiBiT for 1 hour, there was no change in monomeric Tau-HiBiT’s internalization or clearance rate (Figures 5A, 5B and S5A).

**Figure 5.**
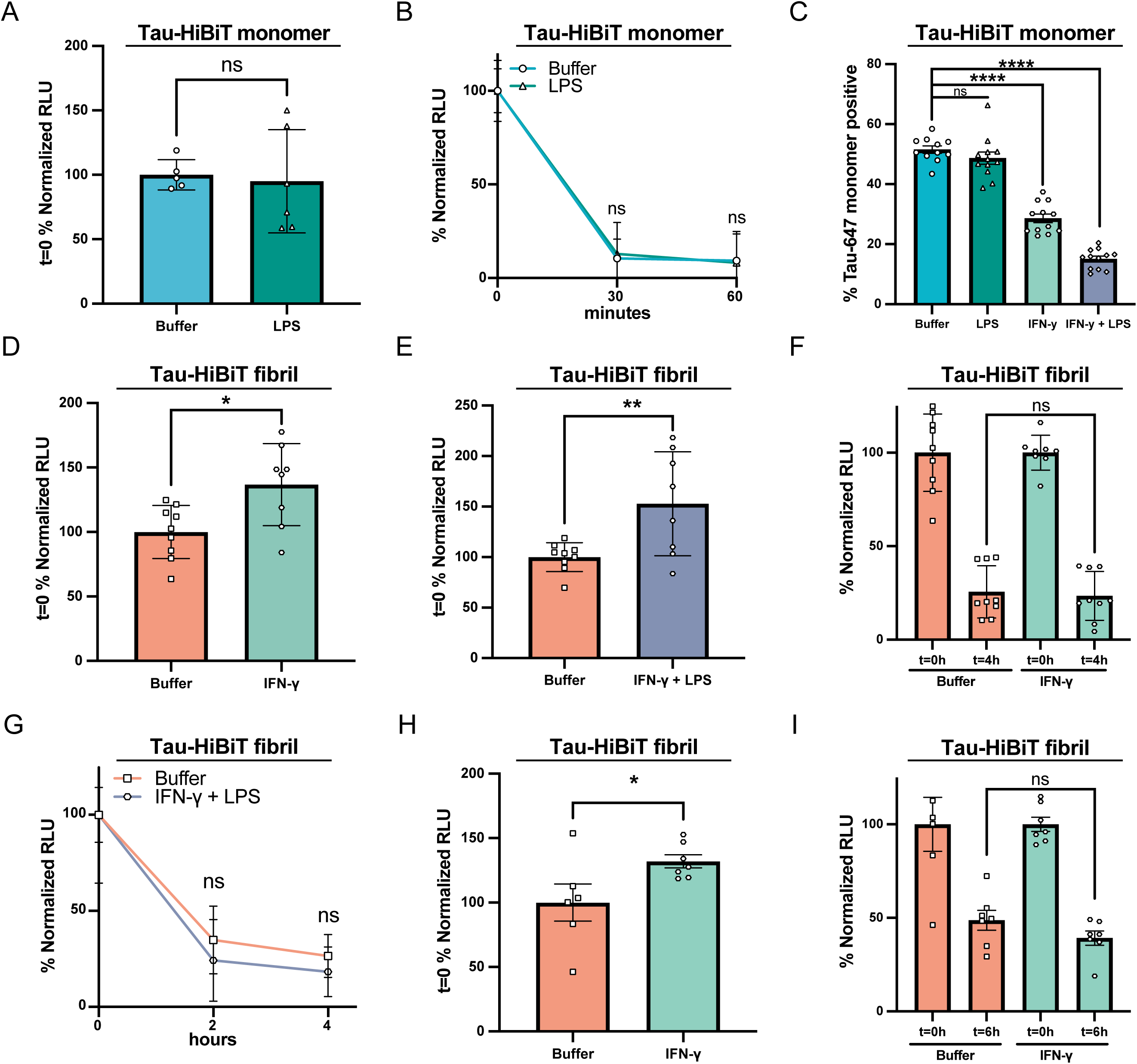
Inflammatory stimuli modulate Tau-HiBiT endocytosis, but do not influence Tau-HiBiT’s clearance rate. (A) Initial uptake (t=0; normalized to Buffer control) of monomeric Tau-HiBiT (100nM) in iTF-Microglia pre-treated with Buffer or LPS (100ng/mL) for 24 h (mean±SD; n ≥ 5 from two independent differentiations; ns = non-significant using unpaired Welch’s t-test). (B) Clearance of monomeric Tau-HiBiT (100nM) in iTF-Microglia pre-treated with Buffer or LPS (100ng/mL) for 24 h. Luminescent signal is normalized to respective initial uptake levels (100%) and cells that were not exposed to Tau-HiBiT (0%) (mean±SD; n ≥ 5 from two independent differentiations; ns = non-significant using a two-way ANOVA; time: *F*(1, 18) = 0.2136, *p* = 0.6495; treatment: *F*(1, 18) = 0.008826, *p* = 0.9262; interaction: *F*(1, 18) = 0.07023, *p* = 0.7940; Sidak’s multiple comparisons test, *p* values indicated on figure, t=0 values are shown for reference but were excluded from statistical analysis). (C) Tau uptake in iTF-Microglia incubated with monomeric Tau-647 (50nM) for 1 h at 37°C after pre-treatment with Buffer, LPS (200ng/mL), IFN-γ (1µg/mL) or LPS + IFN-γ (200ng/mL + 1µg/mL) (mean±SD; n ≥ 11 from three independent differentiations; ns = non-significant, ∗∗∗∗ p < 0. 0001 using a one-way ANOVA, *F*(3, 43) = 134.3, *p* < 0.0001; Tukey’s multiple comparisons test, with p values indicated on figure). (D) Initial uptake (t=0; normalized to Buffer control) of fibrillized Tau-HiBiT (100nM) in iTF-Microglia pre-treated with Buffer or IFN-γ (1µg/mL) for 48 h (mean±SD; n ≥ 8 from three independent differentiations; ∗ p < 0.05 using unpaired Welch’s t-test). (E) Initial uptake (t=0; normalized to Buffer control) of fibrillized Tau-HiBiT (100nM) in iTF-Microglia pre-treated with Buffer or LPS + IFN-γ (100ng/mL + 1µg/mL) for 48 h (mean±SD; n ≥ 8 from three independent differentiations; ∗∗ p < 0.01 using unpaired Welch’s t-test). (F) Clearance of fibrillized Tau-HiBiT (100nM) in iTF-Microglia pre-treated with Buffer or IFN-γ (1µg/mL) for 48 h. Luminescent signal is normalized to respective initial uptake levels (100%) and cells that were not exposed to Tau-HiBiT (0%) (mean±SD; n ≥ 8 from three independent differentiations; ns = non-significant using unpaired Welch’s t-test). (G) Clearance of fibrillized Tau-HiBiT (100nM) in iTF-Microglia pre-treated with Buffer or LPS + IFN-γ (100ng/mL + 1µg/mL) for 48 h. Luminescent signal is normalized to respective initial uptake levels (100%) and cells that were not exposed to Tau-HiBiT (0%) (mean±SD; n = 9 from three independent differentiations; ns = non-significant using a two-way ANOVA; time: *F*(1, 32) = 1.739, *p* = 0.1967; treatment: *F*(1, 32) = 3.068, *p* = 0.0894; interaction: *F*(1, 32) = 0.04798, *p* = 0.8280; Sidak’s multiple comparisons test, *p* values indicated on figure, t=0 values are shown for reference but were excluded from statistical analysis). (H) Initial uptake (t=0; normalized to Buffer control) of fibrillized Tau-HiBiT (100nM) in iMGLs pre-treated with Buffer or IFN-γ (1µg/mL) for 48 h (mean±SD; n ≥ 6 from two independent differentiations; ∗ p < 0.05 using unpaired Welch’s t-test). (I) Clearance of fibrillized Tau-HiBiT (100nM) in iMGLs pre-treated with Buffer or IFN-γ (1µg/mL) for 48 h. Luminescent signal is normalized to respective initial uptake levels (100%) and cells that were not exposed to Tau-HiBiT (0%) (mean±SD; n ≥ 6 from two independent differentiations; ns = non-significant using unpaired Welch’s t-test).

Microglia have also been shown to react towards Interferon Gamma (IFN-γ) and upregulate the expression of inflammatory genes (Hobson et al., 2023; Rock et al., 2005). We therefore compared iTF-Microglia pre-treated with LPS (200ng/mL), IFN-γ (1µg/mL) or LPS + IFN-γ (200ng/mL + 1µg/mL, respectively) and analyzed their ability to endocytose monomeric Tau-647 via flow cytometry, which is a more robust measure of endocytosis. Using this assay, we found that iTF-Microglia pre-treated with LPS had a slight, but insignificant, decrease in tau endocytosis that was substantially enhanced in IFN-γ or LPS + IFN-γ treatment conditions (Figure 5C). We hypothesize that the changes in endocytosis could potentially be due to changes in LRP1 expression with IFN-γ treatment, as has been previously reported (Gorovoy et al., 2010), although we did not explicitly test this. Unfortunately, the reduction in tau endocytosis upon inflammatory treatment was not conducive for performing subsequent tau clearance experiments with monomeric protein.

Based on our prior observations that monomeric and fibrillar tau have diverging endocytic routes, we also wanted to ask how inflammatory stimuli could impact fibrillized tau endocytosis and clearance. Therefore, we pre-treated iTF-Microglia for 48 h with IFN-γ (1µg/mL) and measured tau endocytosis and clearance using the NanoBiT platform. We found that iTF-Microglia pre-treated with IFN-γ had a mild increase in fibrillized Tau-HiBiT endocytosis (Figure 5D), which was further enhanced when iTF-Microglia were pre-treated with both LPS + IFN-γ (Figure 5E). The increase in endocytosis at t=0 was not always consistent between biological replicates/differentiations (Figures S5B and S5C). Additionally, this increase in endocytosis was in stark contrast to our results with monomeric Tau-HiBiT protein, again highlighting potential differences in internalization mechanisms. Despite this higher endocytic rate, we found that the clearance of fibrillized Tau-HiBiT was unchanged upon IFN-γ stimulation (Figure 5F). Clearance was also unchanged with LPS + IFN-γ pre-treatment conditions (Figure 5G) or by stimulating with isolated synaptosomes (Figure S4C). To confirm that this was a consistent phenotype across iPSC-derived microglia models, we also measured fibrillized Tau-HiBiT loss after IFN-γ pre-treatment in the iMGL model. Again, we observed a slight enhancement of fibrillized Tau-HiBiT endocytosis (Figure 5H and S5D), but no change in clearance over time (Figure 5I). With this data we conclude that while inflammatory inducers can enhance endocytosis of fibrillized Tau-HiBiT, they do not, at least under this acute treatment regiment, impact microglial clearance capabilities.

## 4. Discussion

Microglia have a nuanced role in the context of neurodegenerative disease, with multiple lines of experimental evidence to suggest that their canonical “neuro-protective” function is likely compromised in disease. While many studies have shown that microglia can readily internalize tau (Andersson et al., 2019; Bolós et al., 2016; Falkon et al., 2025; Luo et al., 2015), their ability to efficiently process tau during disease has been brought into question. Indeed, some studies have shown that microglia can release tau aggregates back into the media (Brelstaff et al., 2021; Hopp et al., 2018) and removal of microglia from tauopathy mouse models is protective against tau spreading and neuronal loss (Shi et al., 2019; Wang et al., 2022). With the recent advancement of protocols to produce microglia from human iPSCs (Dräger et al., 2022; McQuade et al., 2018), it has now become possible to tease out molecular pathways relevant to human biology that may allow a better understanding of the neurodegenerative disease process in humans.

In this study, we utilized human iPSC-derived microglia to address outstanding questions of how human microglia internalize extracellular tau species and process them. Our results demonstrate that while tau monomers can utilize LRP1 to enter microglia, LRP1 inhibition was less effective on aggregated tau internalization. These results are consistent with prior work showing distinct internalization mechanisms for tau species in human iPSC-derived neurons (Evans et al., 2018), and our prior study where tau monomer uptake was more sensitive to LRP1 inhibition than fibril uptake in H4 neuroglioma cells (Rauch et al., 2020). These results do contrast though with a recent study that showed a minimal influence of LRP1 on the internalization of tau monomer in microglia (Falkon et al., 2025), however this study used primary mouse microglia which may explain the differences in our studies. Taken together, these collective results highlight the importance of cell-type and species when identifying mechanisms that are important for disease-relevant pathways.

To further understand how internalized tau is processed by microglia, we developed a quantitative, live-cell platform using the NanoBiT split-luciferase system, which can measure both total tau endocytosis and the subsequent clearance kinetics of internalized tau proteins. The majority of our mechanistic evidence that HiBiT signal loss reflects endo-lysosomal processing comes from experiments in H4 cells. This choice was driven by technical barriers in iTF-Microglia and iNeuron systems, including acute toxicity of LLOMe and modest CRISPRi knockdown efficiency, which limits reproducibility. While these constraints precluded direct endo-lysosomal manipulations in iPSC-derived cells here, future optimization should confirm whether the same pathways operate in those systems. The NanoBiT system has previously been used to study tau endosomal escape in HEK293 cells and mouse primary neurons (Tuck et al., 2022), and our assay provides a complementary approach for tracking tau endosomal flux. Combining these platforms could allow quantitative tracking of tau fate post-internalization across cell types under both normal and disease conditions. Consistent with known tauopathy pathology, we found that iPSC-derived neurons processed tau far more slowly than microglia, aligning with observations that tau aggregates accumulate in neurons but are rarely found in microglia.

One of the key insights from our study is the observation that internalized monomeric and fibrillar tau are not processed equivalently by cells, including microglia. The use of the NanoBiT system enabled kinetic profiling of extracellular tau handling, revealing that while both monomeric and fibrillized forms of tau are internalized, fibrillar tau exhibits significantly delayed clearance. Although our measurements capture the disappearance of intracellular tau signal, we cannot rule out the possibility that exocytosis contributes to this process, since released tau would no longer be detected in our intracellular assay, and disentangling its role will be an important focus for future work. Overall, our data looking at extracellular tau clearance are consistent with models of intracellular tau degradation, where tau aggregates within cells have been shown to be resistant to proteolytic processing (Guo et al., 2016). Our results add to this growing model of pathogenic tau regulation and highlight that even in cells with high degradative capabilities, such as microglia, tau aggregates can still pose a significant challenge. These findings also promote the idea that the aggregation of tau is a critical switch that shifts tau from a degradable to a more persistent form, ultimately contributing to its pathological accumulation. Prior work in mouse microglial cultures has shown that anti-tau antibodies can be used to enhance uptake and degradation of tau aggregates via FcyR (Andersson et al., 2019). Taken with our results, which point to aggregate state as a critical bottleneck in tau processing, this highlights the potential therapeutic utility of anti-tau antibodies not merely to promote uptake, but to overcome the degradative resistance of fibrillar tau. In the context of our findings, strategies that both facilitate delivery of tau aggregates to lysosomes and enhance lysosomal proteolytic competence may be necessary to achieve meaningful clearance. Such combined approaches could be particularly valuable in disease stages where microglial activation boosts uptake without a parallel increase in degradation efficiency.

Our study also sheds light on the influence of microglial inflammatory activation on tau processing. Acute exposure to IFN-γ, which models classical pro-inflammatory activation, enhanced the uptake of fibrillized tau, suggesting a general increase in phagocytic activity. However, strikingly, this activation did not alter the rate of tau clearance once tau was internalized. This finding decouples microglial activation status from degradative competence, indicating that in the context of acute inflammatory signaling, tau processing is governed by intrinsic properties of the tau species rather than by the cell’s immune polarization. The microglial-immune pathway activation in tauopathy models has been an area of interest for the field, with recent work suggesting that microglia in early disease display a type I interferon signature followed by a phase of partial immune suppression and ultimately a type II interferon response (Rexach et al., 2020). In our study, we modeled acute inflammatory activation by treating microglia with IFN-γ or LPS/IFN-γ. Under these conditions, we observed enhanced uptake of fibrillar tau but no corresponding increase in its clearance. Our data are consistent with the idea that type II interferon signaling in later stages of disease may enhance tau aggregate internalization without necessarily improving degradative capacity and potentially contributing to intracellular accumulation and propagation. The lack of enhanced clearance despite strong inflammatory activation may reflect that lysosomal competence is not intrinsically boosted by this signaling profile, or that fibrillar tau’s inherent resistance to proteolysis remains the limiting step. Together, these findings suggest that the nature of microglial activation, and the balance between uptake and degradation pathways, will be critical variables to consider in therapeutic strategies aimed at enhancing tau clearance.

Human genetics have implicated endo-lysosomal dysfunction as a potential key driver of disease in tauopathies (Van Acker et al., 2019). The mechanistic work provided here further highlights the endo-lysosomal system as a potential therapeutic target for enhancing the clearance of aggregated tau. Strategies aimed at boosting lysosomal capacity or facilitating disaggregation may prove more effective than approaches focused solely on modulating inflammation. Furthermore, our findings suggest that anti-inflammatory interventions alone may be insufficient to restore tau clearance pathways in disease contexts.

An important consideration for our findings is the relevance of iPSC-derived microglia to *in vivo* human biology. While these cells recapitulate many features of primary human microglia, including gene expression profiles and functional behaviors (Hasselmann and Blurton-Jones, 2020), they remain a simplified model. It is known that microglia cultured *in vitro* express a basal expression of inflammatory proteins which may be a confounding factor when attempting to study the influence of inflammation on Tau-HiBiT’s degradation (Cadiz et al., 2022; Lloyd et al., 2024; Warden et al., 2023). Additionally, our studies modeled an acute inflammatory condition which contrasts with chronic diseases such as Alzheimer’s disease. It is also known that iPSC reprogramming resets aging markers (Mahmoudi and Brunet, 2012), therefore by utilizing iPSC-derived microglia, our work in modeling inflammatory conditions may not fully recapitulate the complexity of the aging neuroinflammation environment within the CNS. Future studies that integrate iPSC-microglia into 3D brain organoids or chimeric mouse models may provide additional insights into how tau handling occurs in a more physiological environment, particularly in the context of chronic disease and aging.

## 5. Conclusions

In conclusion, our study delineates the distinct pathways through which human microglia process different forms of tau and positions the lysosomal degradation step as the key limiting factor for clearance of aggregated tau. By leveraging a scalable, quantitative approach, we offer a new tool for dissecting tau–microglia interactions and identifying novel therapeutic strategies. These findings contribute to a growing understanding of microglial contributions to tauopathy and emphasize the need to consider aggregation state when evaluating mechanisms of tau clearance in neurodegenerative disease.

## Supporting information

Supplemental Western Blots

## CRediT authorship contribution statement

**Andrew Shultz:** Conceptualization, Methodology, Formal analysis, Investigation, Writing – original draft, Writing – review & editing, Visualization, Funding Acquisition. **Anna Vincze:** Investigation. **Todd T. Yau:** Investigation. **Emma L. Poirier:** Investigation. **Hannah Demanche:** Investigation. **Jennifer N. Rauch:** Conceptualization, Methodology, Formal analysis, Writing – original draft, Writing – review & editing, Visualization, Supervision, Funding Acquisition.

## Declaration of competing interest

The authors declare no competing interests.

## Acknowledgements

This work was supported by the National Institutes of Health (R00AG064116, R01AG077672) to J.N.R. This research was partially supported by National Research Service Award T32 GM139789 from the National Institutes of Health awarded to A.S. The funders had no role in study design, data collection and analysis, decision to publish, or preparation of the manuscript. We thank Martin Kampmann for the use of the iPSC lines used to generate the iTF-Microglia and iNeurons. We are thankful to the University of Massachusetts Core Facilities for use of their Biophysical Characterization Core (RRID:SCR_022357) and Electron Microscopy Core. The graphical abstract and some figure images were created using Biorender.

## Figure Captions

**Supplemental Figure 1.**

(A) Representative ICC of iTF-Microglia after differentiation stained for TMEM119 (Green). Nuclei are stained with Hoechst (Blue); scale bar = 50µm.

(B) RT-qPCR of iTF-Microglia after differentiation. Relative gene expression was calculated using the delta Ct (ΔΔCt) method with GAPDH used as a housekeeping gene; n=3.

(C) HEK-Blue hTLR4 cells were incubated with endotoxin standards or recombinantly expressed proteins. Conditioned media was collected and quantified according to manufacturer’s recommendations. Calculated OD(620nm) for 0.1 EU/mL and 0.05 EU/mL are indicated with dashed-lines (mean±SD; n = 3).

(D) RT-qPCR of LRP1 gene expression in iTF-Microglia with (TMP, 20µM) or without (DMSO) LRP1 knockdown. Relative gene expression was calculated using the delta Ct (ΔΔCt) method with GAPDH used as a housekeeping gene (mean±SD; n = 3 from one independent day).

(E) Western Blot of iTF-Microglia with (TMP, 20µM) or without (DMSO) LRP1 knockdown.

(F) Transferrin uptake in iTF-Microglia with (TMP, 20µM) or without (DMSO) LRP1 knockdown (mean±SD; n = 3).

(G) Tau uptake in iMGLs incubated with monomeric Tau-647 (50nM) or Tau-647 (50nM) + RAP (250nM) for 1 h at 37°C (mean±SD; n = 3; ∗∗p < 0. 01 using unpaired Welch’s t-test)

(H) TEM negative stain image of Tau-HiBit fibrils. Scale bar = 200nm.

(I) TEM negative stain image of Tau-HiBiT monomer. Scale bar = 200nm.

(J) ThioflavinT fluorescence measured during aggregation of Tau-HiBiT (mean±SD; n = 9).

(K) Representative flow cytometry graph from iTF-Microglia after Tau-647 monomer uptake, corresponding to data in Figure 1C.

(L) Representative flow cytometry graph from iTF-Microglia after Tau-647 fibril uptake, corresponding to data in Figure 1E.

**Supplemental Figure 2.**

(A) RT-qPCR of RAB7A gene expression in H4i cells. Relative gene expression was calculated using the delta Ct (ΔΔCt) with GAPDH used as a housekeeping gene (mean±SD; n = 3).

(B) Western Blot of H4i cells expressing NTC sgRNA and RAB7A sgRNA.

**Supplemental Figure 3.**

(A) Western Blot of pTau-HiBiT protein. PHF-1 antibody recognizes pS396/pS404.

(B) Luminescence of Tau-HiBiT proteins (100nM to 10pM) (mean±SD; n = 2).

**Supplemental Figure 4.**

(A) Quantification of cytokine array panel shown in (B).

(B) Representative cytokine array panel. Membranes from the Proteome profiler human cytokine array kit were incubated with conditioned media from iTF-Microglia that were stimulated with Buffer (Top) or LPS (100ng/mL, 24 hours) (Bottom).

(C) Clearance of fibrillized Tau-HiBiT (50nM) in iTF-Microglia treated with rat synaptosomes (0.5mg/mL) for 24 h (mean±SD; n = 3).

**Supplemental Figure 5.**

(A) Raw t=0 RLU data corresponding to normalized data presented in Figure 5A.

(B) Raw t=0 RLU data corresponding to normalized data presented in Figure 5D.

(C) Raw t=0 RLU data corresponding to normalized data presented in Figure 5E.

(D) Raw t=0 RLU data corresponding to normalized data presented in Figure 5H.

For all data, biological replicates from independent differentiations are shown in differing colors and p values from unpaired Welch’s t-test for each cohort is displayed.

**Supplemental Uncropped Western blots.**

Images provided of uncropped western blots for Figure S1, Figure S2B, and Figure S3A.

## Notes

### Competing Interest Statement

The authors have declared no competing interest.

### Summary of Updates

We have added additional replicates and differentiations to strengthen the rigor of our findings, updated the figure layout and color scheme for improved clarity, included graph titles to aid interpretation, adjusted our statistical analyses where appropriate, tempered our language to accurately describe our experimental design, and expanded the discussion to better highlight the significance of our results.

