## Supplementary figures and images for "Extracellular tau clearance is governed by its aggregation state and independent of microglial activation by LPS and IFN-γ"

### Figure S1

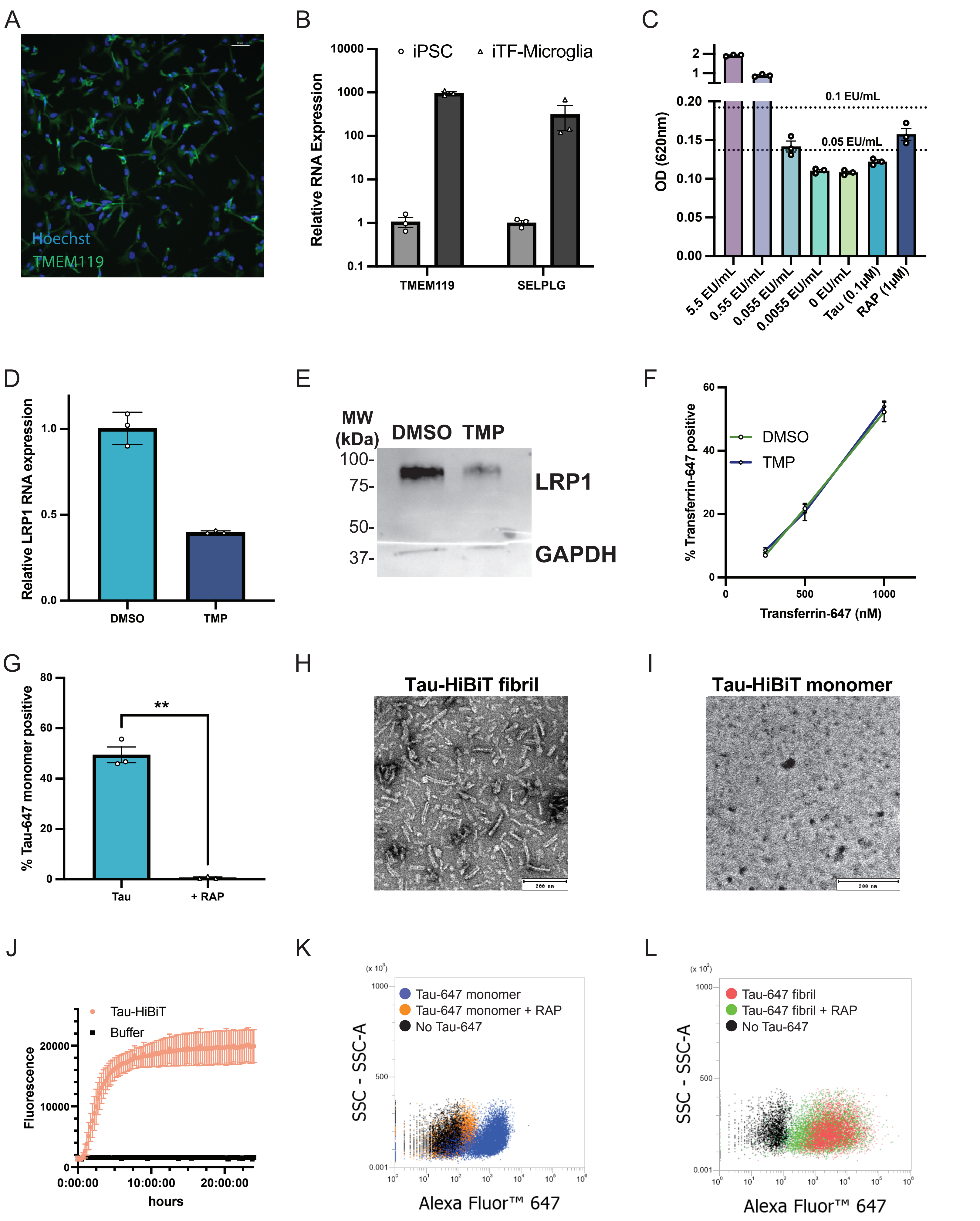

### Figure S2

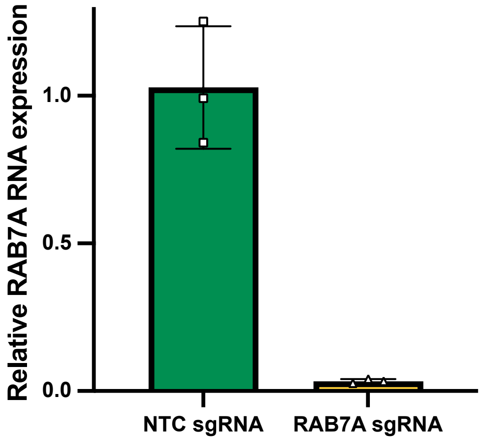

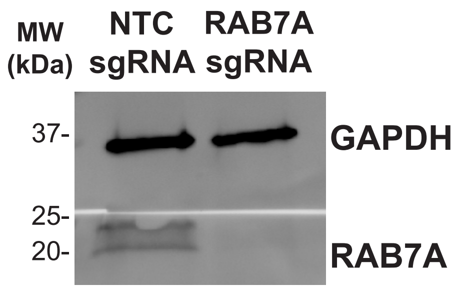

### Figure S3

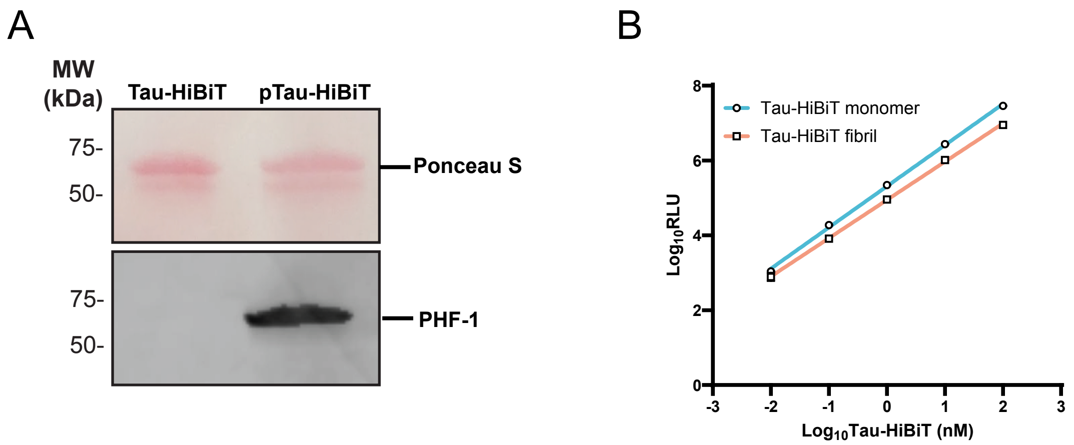

### Figure S4

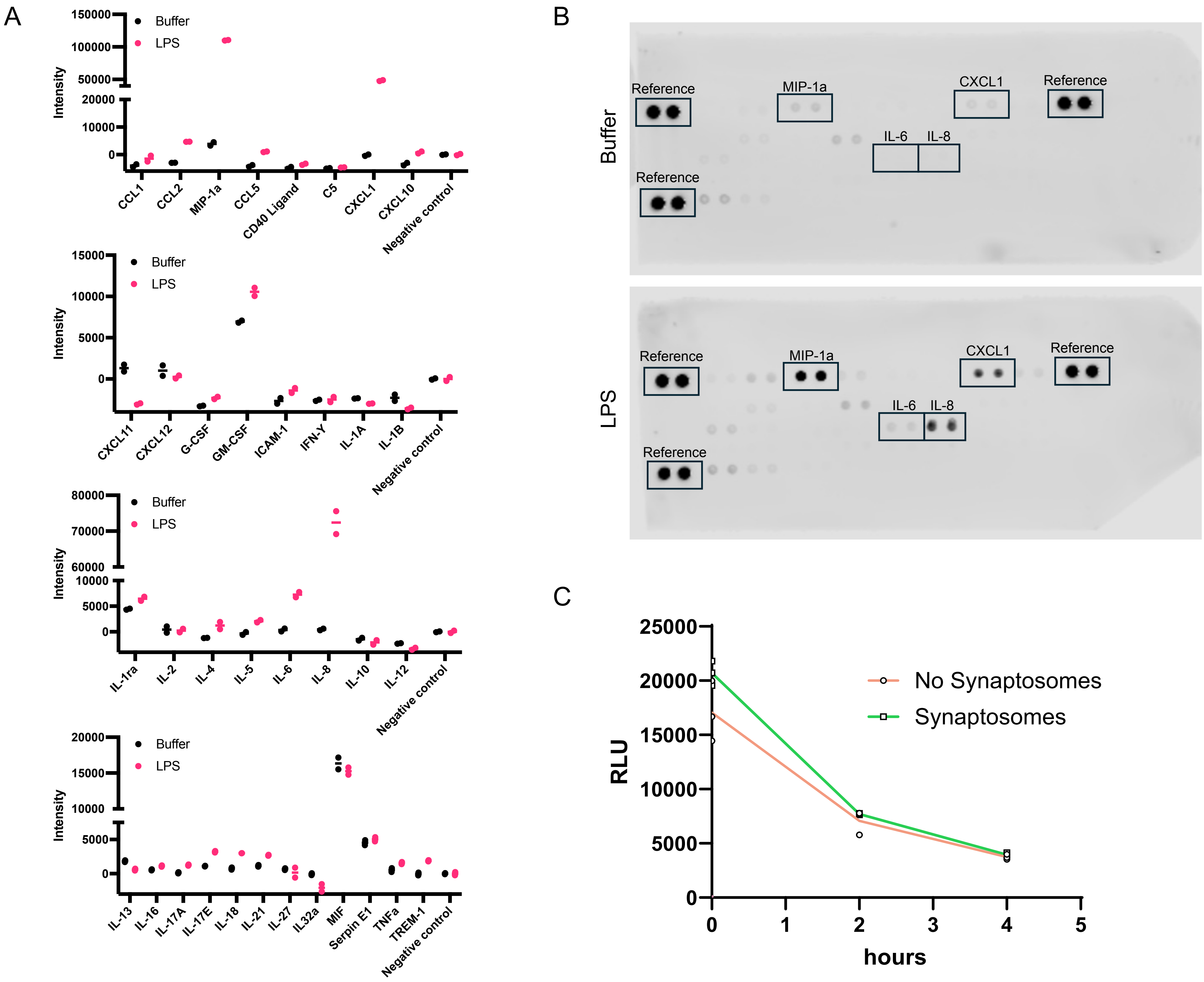

### Figure S5

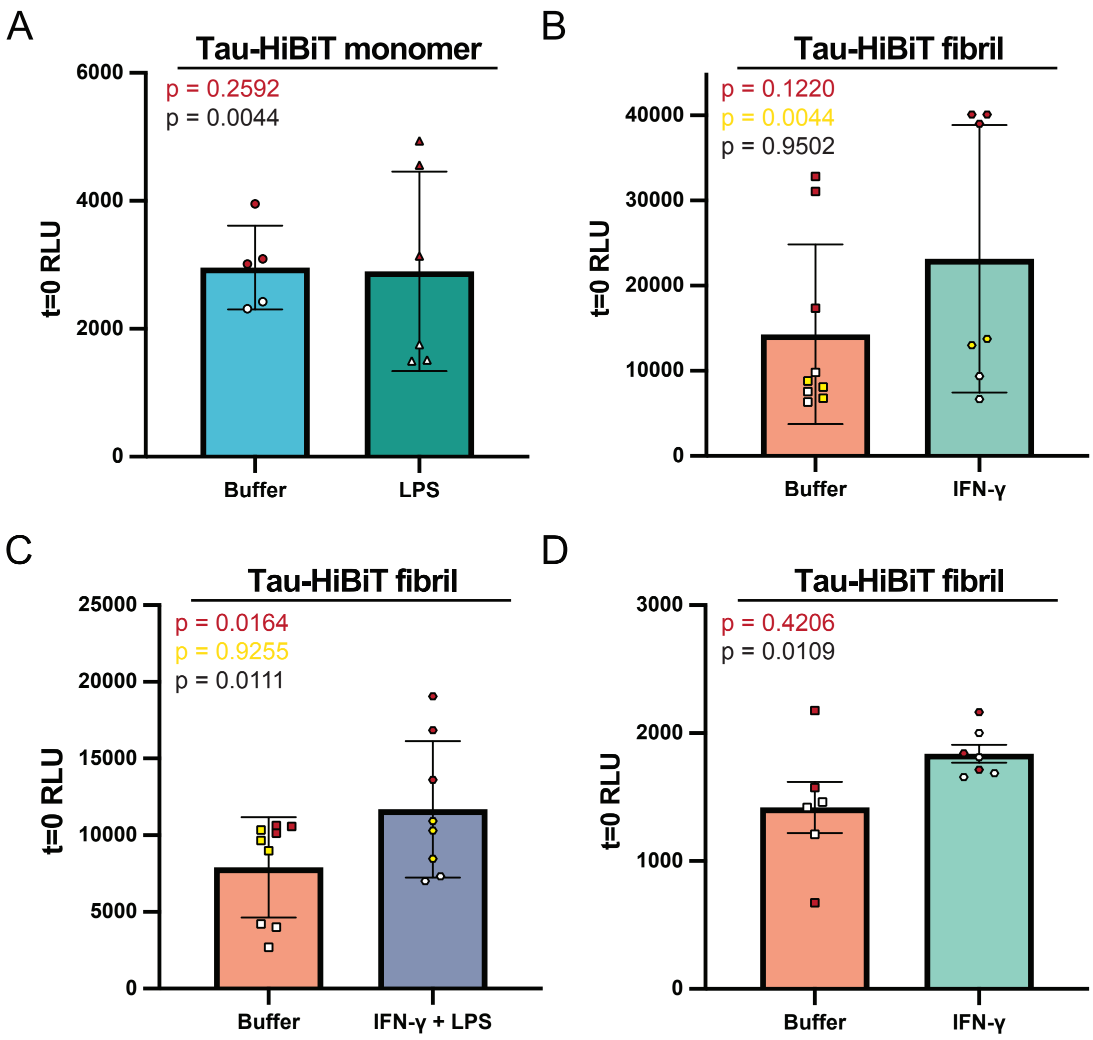

### Supplemental Western Blots

#### Figure S1E

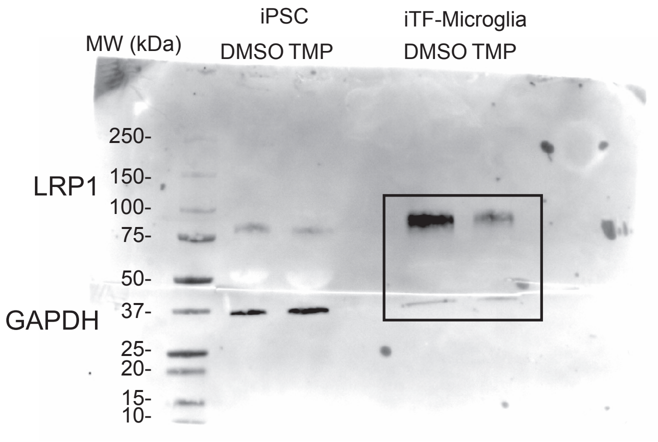


#### Figure S1E

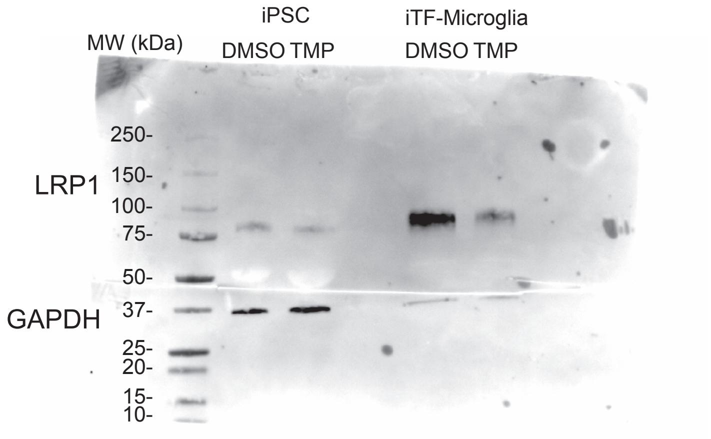


#### Figure S2A

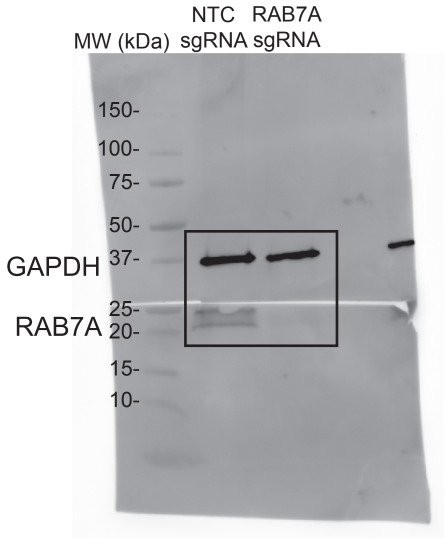


#### Figure S2A

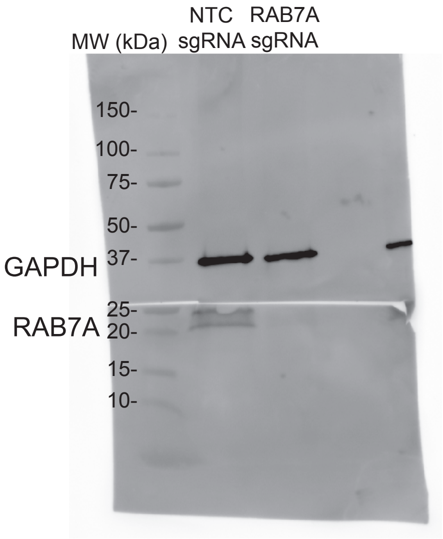


#### Figure S3A

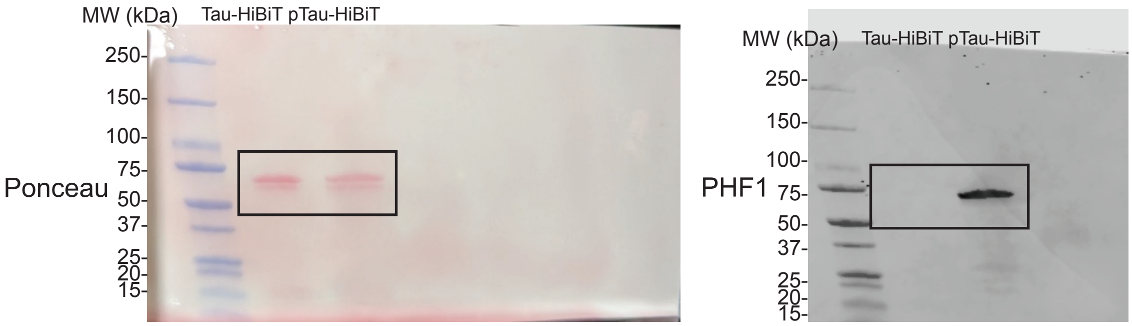


#### Figure S3A

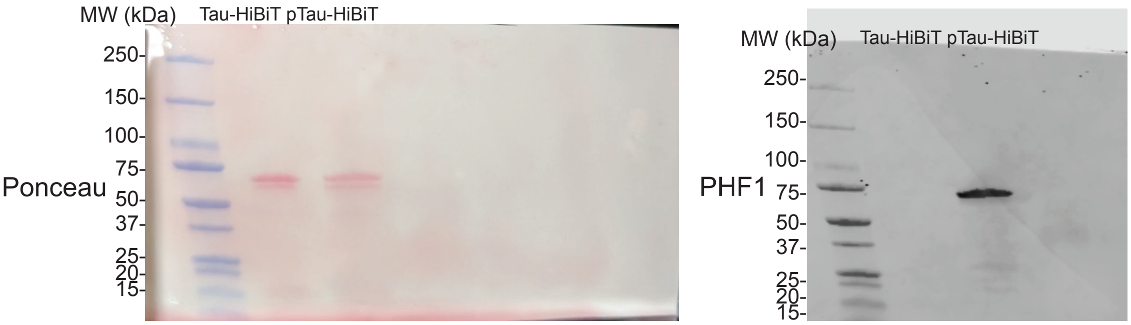
